# Cellular acidosis triggers MondoA transcriptional activity by driving mitochondrial ATP production

**DOI:** 10.1101/376368

**Authors:** Blake R. Wilde, Zhizhou Ye, Donald E. Ayer

## Abstract

MondoA and its transcriptional target thioredoxin-interacting protein (TXNIP) constitute a regulatory loop that senses glycolytic flux and controls glucose availability. Cellular stress also triggers MondoA activity and TXNIP expression. To understand how MondoA integrates glucose and stress signals, we studied its activation by acidosis. We found that acidosis drives mitochondrial ATP (mtATP) synthesis. The subsequent export of mtATP from mitochondria via adenine-nucleotide transporter and voltage-dependent anion channel, and the enzymatic activity of mitochondria-bound hexokinase results in the production of glucose-6-phosphate (G6P), a known activator of MondoA transcriptional activity. MondoA localizes to the outer-mitochondrial membrane (OMM), and in response to G6P, shuttles to the nucleus and activates transcription. Our data suggests that MondoA is a required feature of a glucose- and mtATP-dependent, OMM-localized signaling center. We propose MondoA functions as a coincidence detector and its ability to sense glucose and cellular stress is coupled to the concerted production of G6P.

## INTRODUCTION

Glucose is a major source of carbons for the production of ATP and biosynthetic intermediates. Dysregulation of glucose uptake and metabolism underlies many diseases including cancer and diabetes (Petersen et al., 2017, Hay, 2016). Thus, it is important to understand the precise molecular mechanisms that regulate glucose homeostasis in normal and pathological settings.

The paralogous transcription factors MondoA and ChREBP (MondoB) are sentinel regulators of glucose-induced transcription and their activity is highly, if not entirely, dependent on glucose (Stoltzman et al., 2008, Richards et al., 2017, Peterson et al., 2010, Stoltzman et al., 2011, Ma et al., 2005). Work by our lab and others has established glucose-6-phosphate (G6P) as a key regulatory signal that drives Mondo transcriptional activity (Stoltzman et al., 2008, Li et al., 2010). Other hexose-6-phosphates, fructose-2,6-bisphosphate, and xylulose-5-phosphate are also thought to drive Mondo-dependent transcription, yet the molecular mechanisms are not well-defined (Kabashima et al., 2003, Petrie et al., 2013, Stoltzman et al., 2011).

MondoA controls the glucose-dependent expression of thioredoxin-interacting protein (TXNIP), which has a number of critical cellular functions (Anderson, 2016, Shalev, 2014, O’Shea and Ayer, 2013). The best characterized among these is as a suppressor of glucose uptake (Stoltzman et al., 2008, Wu et al., 2013, Hui et al., 2008). Thus, MondoA and TXNIP – the MondoA/TXNIP axis – make up a negative feedback loop that maintains cellular glucose homeostasis. High TXNIP is anti-correlated with glucose uptake in human tumors and is a predictor of better overall survival in cancer patients, establishing the MondoA/TXNIP axis as an important prognostic factor in cancer (Lim et al., 2012, Chen et al., 2010, Shen et al., 2015).

MondoA shuttles from the outer mitochondrial membrane (OMM) to the nucleus where it drives transcriptional circuits that control cellular fuel choice (Billin et al., 2000, Sans et al., 2006, Stoltzman et al., 2008). In addition to being regulated by glucose, a functional electron transport chain (ETC) is also required for MondoA-dependent transcription (Yu et al., 2010, Han and Ayer, 2013), yet the ETC-derived signal remains unknown. It is also unclear how glycolytic and mitochondrial signals converge to regulate MondoA transcriptional activity. Nevertheless, because MondoA responds to both glycolysis and mitochondrial respiration, MondoA may function as a master sensor of cellular energy charge.

TXNIP expression is driven by a number of cellular stresses. For example, serum starvation, lactic acidosis/low pH, ultraviolet and gamma irradiation, endoplasmic-reticulum stress and microgravity (Elgort et al., 2010, Chen et al., 2010, Junn et al., 2000, Versari et al., 2013, Oslowski et al., 2012). However, little is known about how TXNIP expression is regulated by this diverse collection of signals. TXNIP expression is highly, if not entirely, dependent on MondoA and glucose, suggesting that at least some of these stresses may impact MondoA activity and/or the availability of glucose-derived metabolites.

Intracellular acidification is a metabolic stress intrinsic to proliferative cells that results from increased glycolytic flux and consequent lactate production. Cancer cells initiate a homeostatic response to intracellular acidification to restore physiological pH that includes export of lactate, slowing of glycolysis and restricting glucose uptake (Webb et al., 2011, Gunnink et al., 2014). pH-regulation of glycolytic flux and proton transport have been well-studied (Webb et al., 2011), and our previous work suggests a role for the MondoA/TXNIP axis in normalizing cellular pH. For example, lactic acidosis triggers MondoA-dependent TXNIP expression and decreased glucose uptake (Chen et al., 2010). This suppression of glucose uptake requires both MondoA and TXNIP, yet how lactic acidosis activates MondoA transcriptional activity was not investigated.

Here we show that acidic pH drives MondoA transcriptional activity by increasing mitochondrial ATP (mtATP) synthesis. mtATP is used by mitochondria-bound hexokinase to generate G6P from cytoplasmic glucose, which subsequently drives MondoA nuclear accumulation and transcriptional activity. These results suggest a critical role for the MondoA/TXNIP axis in coordinating the transcriptional and metabolic response to the cell’s principal energy sources, glucose and mtATP, and in maintaining energy homeostasis in response to nutrient hyper-abundance.

## RESULTS

### Low pH medium drives MondoA-dependent TXNIP expression

We previously showed that lactic acidosis triggers the MondoA/TXNIP axis (Chen et al., 2010). This finding raised the intriguing possibility that intracellular pH modulates MondoA transcriptional activity. Proton export is primarily regulated by the monocarboxylate transporters (MCTs) and sodium-hydrogen antiporter 1 (NHE1)(Webb et al., 2011). We used publicly available gene expression data to correlate TXNIP expression with MCTs and NHE1. TXNIP expression is inversely correlated with MCT4 in breast cancer, MCT1 in lung cancer and NHE1 in brain cancer (Figure 1A). TXNIP expression was also anti-correlated with MCTs and NHE1 in non-transformed tissues (Figure 1 – figure supplement 1A-B). Further, we identified a correlation between TXNIP expression and an acidosis gene-signature in breast cancer (Figure 1B). These data suggest that intracellular pH per se, rather than a lactic acidosis-dependent signaling event, controls MondoA transcriptional activity.

**Figure 1.**
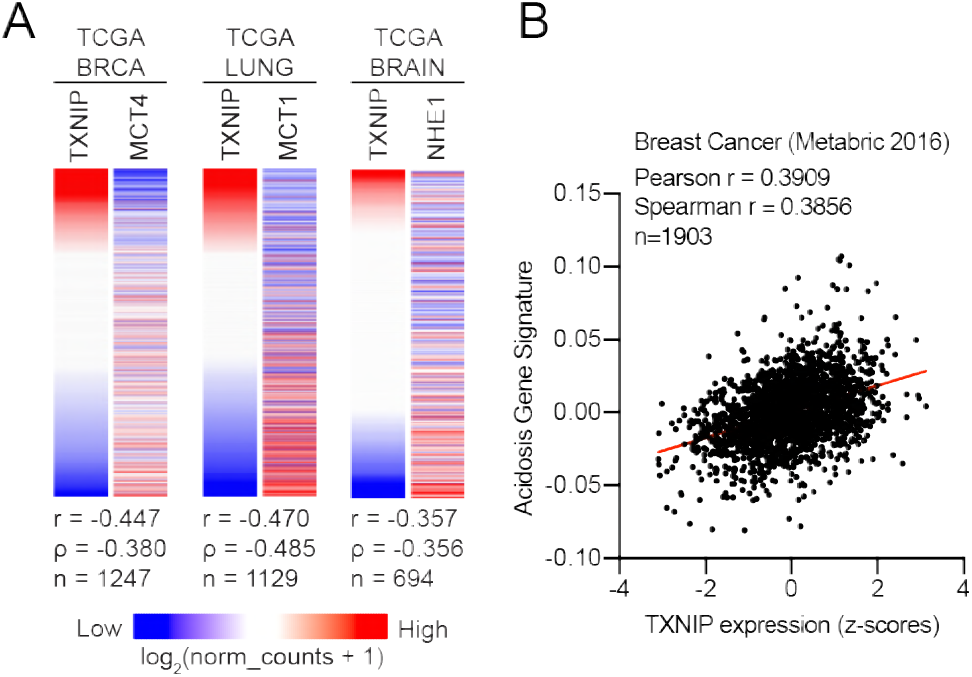
TXNIP correlates with genes that regulate intracellular pH. (**A**) Heatmaps depicting the expression of TXNIP mRNA compared to MCT4 (breast cancer), MCT1 (lung cancer) and NHE1 (brain cancer). All expression data was collected from TCGA. Spearman and Pearson correlation statistics are reported as r and ρ, respectively. (**B**) An acidosis gene signature was determined for the 2016 METABRIC breast cancer dataset. These scores were compared to TXNIP expression from the dataset and correlation statistics were performed.

To better understand the effects of acidosis on MondoA transcriptional activity, we treated cells with Hank’s balanced salt solution (HBSS), which mimics the nutrient-poor extracellular environment cancer cells experience *in vivo*. HBSS has minimal pH-buffering capacity and in 5% CO2 has an acidic pH of ~6.4. HBSS treatment of mouse embryonic fibroblasts (MEFs) increased TXNIP mRNA and protein expression, and decreased glucose uptake (Figure 2A-B, Figure 2 – figure supplement 1A). HBSS is weakly buffered due to its low level of sodium bicarbonate (0.35 g/L). Supplementing HBSS with sodium bicarbonate to 3.7 g/L raised the pH to 7.5 and prevented TXNIP induction (Figure 2C, Figure 2 – figure supplement 1B). Conversely, decreasing sodium bicarbonate in DMEM to 0.37 g/L decreased the pH to ~6.5 and induced TXNIP expression (Figure 2D, Figure 2 – figure supplement 1C). HBSS and DMEM with low sodium-bicarbonate (DMEM^Acidic^) were used throughout this study to mimic extracellular acidification. To determine whether TXNIP induction is mediated by sodium bicarbonate or pH, we increased the pH of HBSS and DMEM^Acidic^ to 7.4. This prevented TXNIP induction (Figure 2 – figure supplement 1C), confirming that low pH rather than low sodium bicarbonate is primarily responsible for HBSS-and DMEM^Acidic^-driven MondoA transcriptional activity.

**Figure 2.**
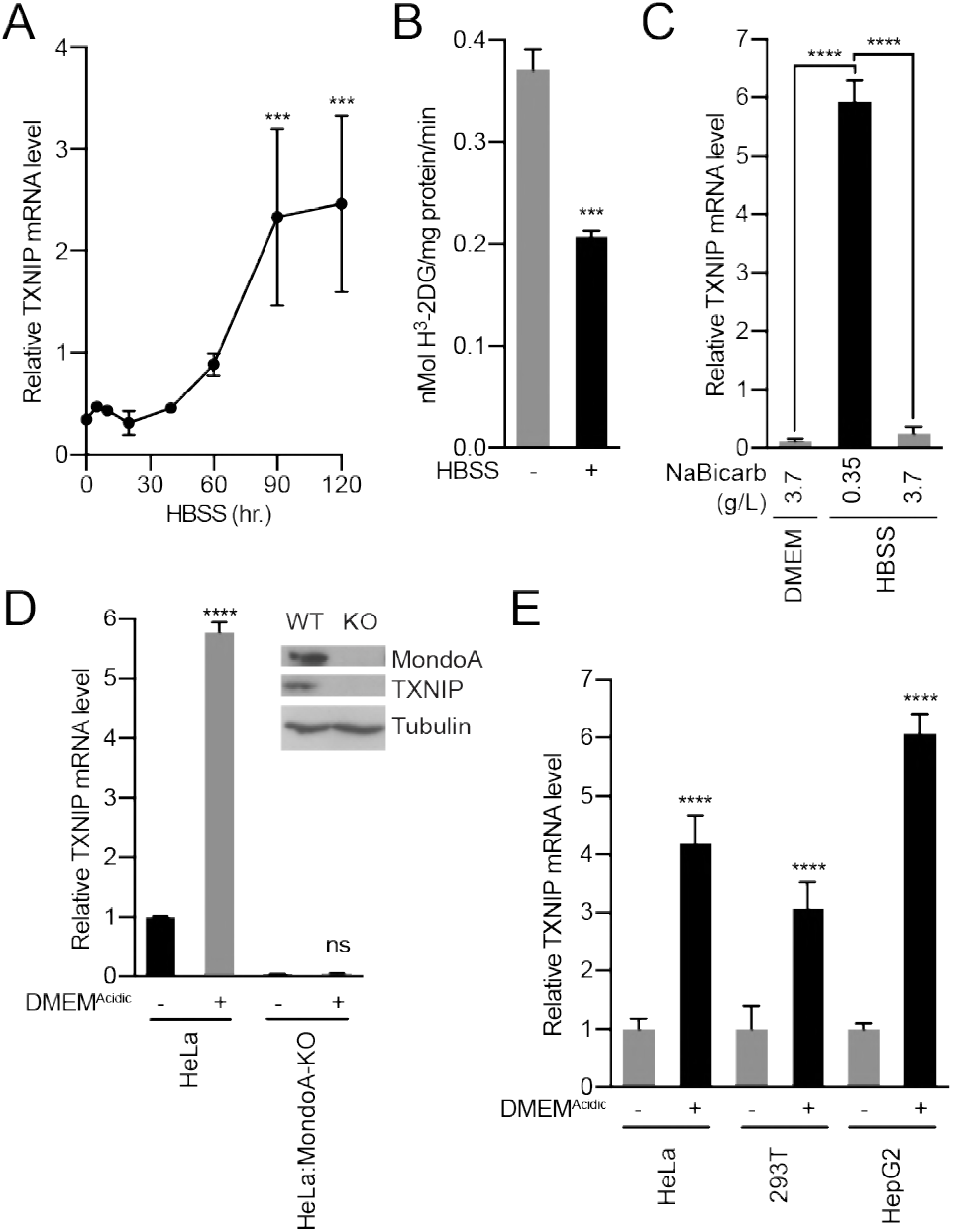
Acidosis drives MondoA transcriptional activity. (**A**) Mouse embryonic fibroblasts (MEFs) treated with HBSS for the indicated amounts of time and TXNIP mRNA levels were determined by reverse transcriptase-quantitative PCR (RT-qPCR). (**B**) Glucose uptake was determined by quantifying the rate of H^3^-2-deoxyglucose uptake in MEFs treated with HBSS. (**C**) TXNIP mRNA levels from MEFs treated with DMEM, HBSS and HBSS supplemented with sodium bicarbonate to the same level as in DMEM (3.7 g/L). (**D**) CRISPR/Cas9 was used to disrupt the expression of MondoA in HeLa cells. Immunoblot of HeLa and HeLa:MondoA-KO cells. Consistent with our previous findings, loss of MondoA prevented TXNIP expression. TXNIP mRNA levels from HeLa and HeLa:MondoA-KO cells treated with DMEM^Acidic^. (**E**) TXNIP mRNA levels in HEK-293T, HeLa and HepG2 cells treated with DMEM^Acidic^.

MondoA is necessary and sufficient for TXNIP induction (Stoltzman et al., 2011, Stoltzman et al., 2008). Consistent with this, HBSS increased TXNIP expression in MondoA^+/+^ MEFs but not in MondoA^-/-^ MEFs (Figure 2 – figure supplement 2A). Reconstituting MondoA^-/-^ MEFs with MondoA rescued TXNIP induction (Figure 2 – figure supplement 2A). Further, TXNIP was induced in HeLa cells treated DMEM^Acidic^ but not in HeLa cells with disrupted MondoA expression (HeLa:MondoA-KO cells, Figure 2D). Finally, DMEM^Acidic^ induced TXNIP expression in three cell lines of different lineages: HeLa, HepG2 and 293T cells (Figure 2E).

We next determined the effects of acidosis on MondoA transcriptional activity. Heterodimerization with Mlx is required for MondoA nuclear translocation and binding to carbohydrate responsive elements (ChoREs) in the promoters of its target genes (Stoltzman et al., 2011, Peterson et al., 2010, Minn et al., 2005, Stoltzman et al., 2008). MondoA(I766P), which does not interact with Mlx (Stoltzman et al., 2008), was unable to rescue TXNIP induction in MondoA^-/-^ MEFs (Figure 2 – figure supplement 2A), indicating a requirement for the MondoA:Mlx heterocomplex. Further, HBSS induced the activity of a TXNIP-promoter luciferase reporter, but not when the ChoRE sequence was mutated (Figure 2 – figure supplement 2B). Finally, HBSS treatment led to increased MondoA occupancy at the TXNIP promoter (Figure 2 – figure supplement 2C). Together these data establish that acidosis drives MondoA transcriptional activity.

### MondoA is required for the transcriptional response to acidosis

To determine the contribution of MondoA to acidosis-driven gene expression we conducted RNA-sequencing on mRNA from HeLa and HeLa:MondoA-KO cells treated with DMEM^Acidic^ for 4 hours. Using a 1.5-fold cut off and an adjusted p-value of ≤ 0.01, we identified 617 differentially regulated genes in HeLa cells treated with DMEM^Acidic^. Of these, 227 were not regulated in HeLa:MondoA-KO cells, suggesting that MondoA contributes to nearly 37% of the acidosis-driven transcriptional response. We next used regression analysis to look for genes that are affected by both DMEM^Acidic^ treatment and genotype. Loss of MondoA prevented the induction/suppression of several acidosis-regulated genes; however, only two genes, TXNIP and one of its paralogues, ARRDC4, were entirely dependent on MondoA (Figure 3A).

**Figure 3.**
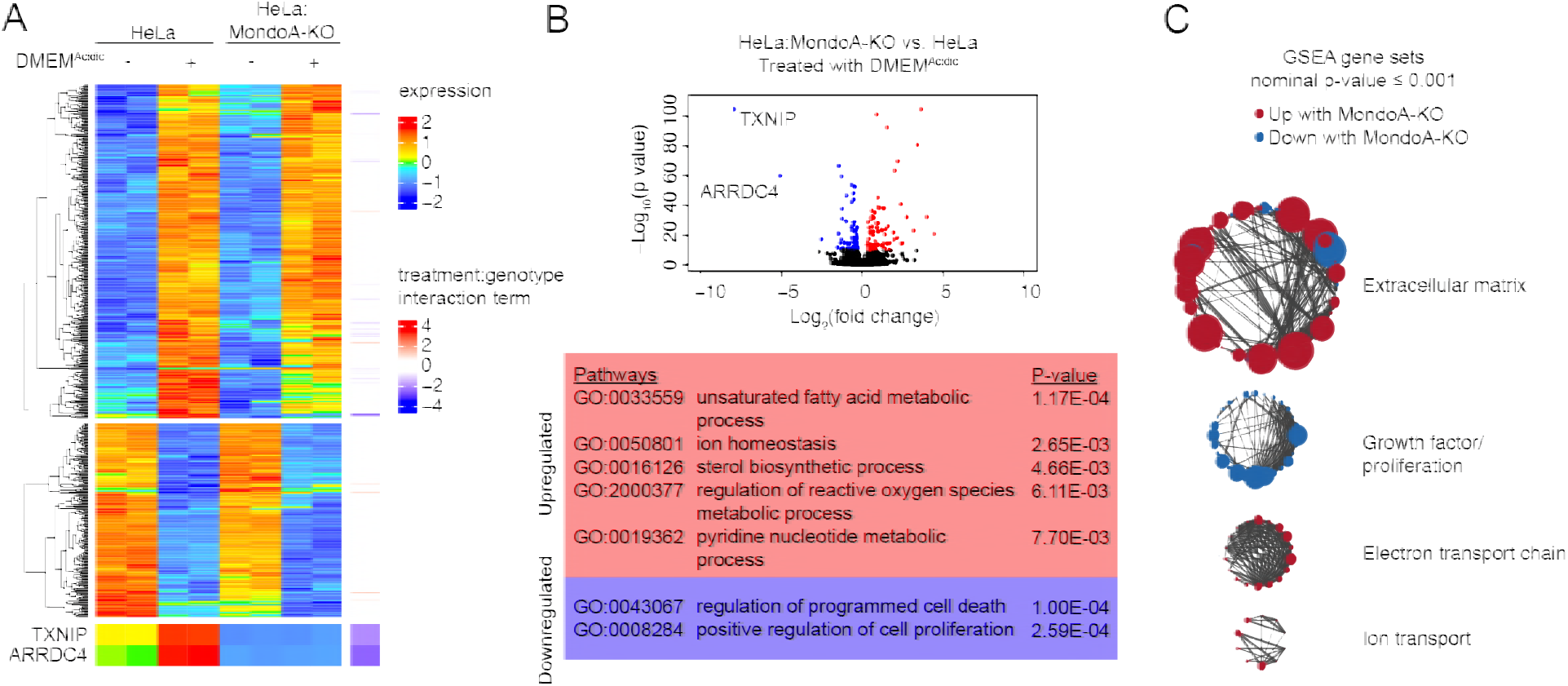
The MondoA-dependent acidosis response. RNA-sequencing was used to determine differentially regulated genes for HeLa and HeLa:MondoA-knockout cells treated with DMEM^Acidic^ for four hours. Differentially regulated genes were determined. (**A**) Heatmaps depicting TXNIP, ARRDC4 and the top 500 differentially regulated genes in HeLa cells treated with DMEM^Acidic^. The genotype:treatment interaction term was calculated using DESeq2 and indicates the influence of both genotype and treatment on differential expression. (**B**) Volcano plot of log_2_(fold-change) of HeLa cells treated with DMEM^Acidic^ compared to HeLa:MondoA-KO cells treated with DMEM^Acidic^. Genes with an adjusted p-value ≤ 1E-10 that are upregulated or downregulated in HeLa:MondoA-KO cells are indicated in red and blue, respectively. Overrepresentation analysis was performed for the upregulated and downregulated genes. Enriched pathways and their respective p-values are given in the red and blue boxes for upregulated and downregulated genes, respectively. (**C**) GSEA and leading edge analysis was conducted for HeLa cells treated with DMEM^Acidic^ compared to HeLa:MondoA-KO cells treated with DMEM^Acidic^. Depicted are networks of gene sets with a nominal p-value ≤ 0.001. Node colors are representative of whether the gene set was positively (red) or negatively (blue) enriched. Node size represents gene set size. Connecting line thickness represents similarity between two nodes.

We next performed pathway analysis on genes differentially regulated in HeLa and HeLa:MondoA-KO cells treated with DMEM^Acidic^. Consistent with the results above, TXNIP and ARRDC4 were the most highly MondoA-dependent genes, with log_2_(fold-changes) of 7.9 and 5.1, respectively (Figure 3B). We identified 157 other differentially regulated genes in HeLa:MondoA-KO cells (adjusted p-value ≤ 1E-10). Pathways that were upregulated in HeLa:MondoA-KO cells were enriched for fatty acid metabolism, sterol biosynthesis, ion homeostasis, ROS metabolism, and pyridine metabolism pathways, whereas cell death and proliferation pathways were downregulated (Figure 3B, Figure 3 – table supplement 1). Further, we conducted gene set enrichment analysis (GSEA) on HeLa and HeLa:MondoA-KO cells treated with DMEM^Acidic^ using all pathways in the Molecular Signatures Database. We identified 588 gene sets that were enriched with a nominal p-value < 0.001 (Figure 3 – table supplement 2). Leading edge analysis highlighted extracellular matrix remodeling, electron transport chain and ion transport as upregulated, and growth-factor/proliferation as downregulated in HeLa:MondoA-KO cells (Figure 3C). Together these data show that MondoA is required for the transcriptional response to DMEM^Acidic^ treatment and suggests that MondoA may have an essential role in an adaptive response to acidosis.

### MondoA is dependent upon mitochondrial ATP

Given the predominant role of MondoA in the transcriptional response to acidosis, we sought to determine how acidic pH triggers MondoA transcriptional activity. Previous reports show that treating cells with low pH medium drives intracellular acidification (Adams et al., 2006, Wahl et al., 2000). In an effort to determine the intracellular site of action of low pH on MondoA activity, we used compartment-selective ionophores to alter proton concentrations in various cellular compartments. Monensin, which drives cytosolic alkalization, abrogated HBSS-induced TXNIP expression (Figure 4 – figure supplement 1A). By contrast, chloroquine which disrupts acidification of endosomes/lysosomes, had no effect on TXNIP induction (Figure 4 – figure supplement 1A). Finally, the mitochondrial ionophore FCCP, prevented HBSS-driven TXNIP expression (Figure 4 – figure supplement 1B). Together these results suggest that cytosolic and/or mitochondrial proton gradients, but not pH-dependent changes in the endosome/lysosome, are critical for the activation of the MondoA/TXNIP axis.

**Figure 4.**
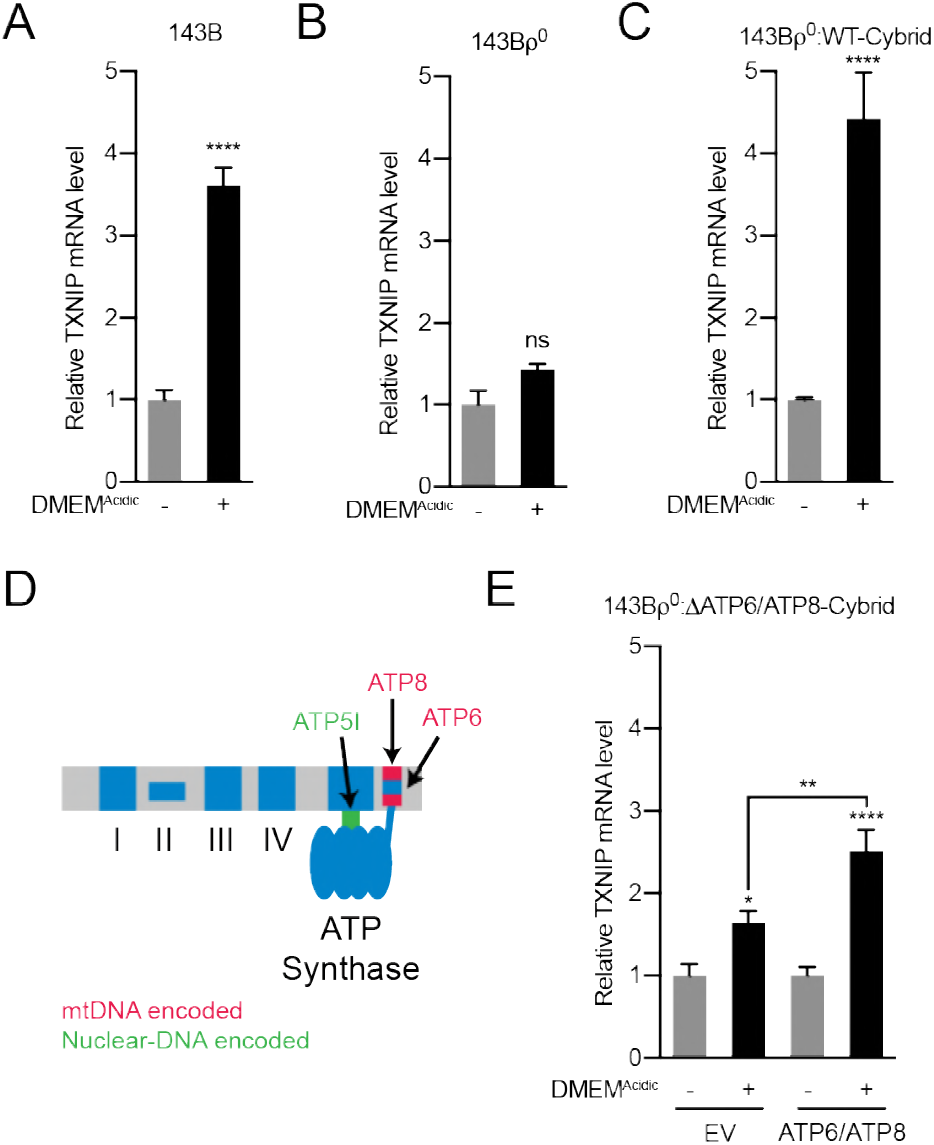
Acidosis-driven MondoA transcriptional activity requires mitochondrial ATP production. TXNIP mRNA level following treatment with DMEM^Acidic^ in (**A**) 143B osteosarcoma cells, (**B**) 143Bρ^0^ cells which lack mtDNA, and (**C**) 143Bρ^0^:WT-Cybrid cells which have restored wild type mitochondria. (**D**) Schematic depicting nuclear- and mitochondrial-DNA encoded components of the ETC. (**E**) TXNIP mRNA level following treatment with DMEM^Acidic^ in 143Bρ^0^:ΔATP6/ATP8-Cybrid cells expressing empty vector or nuclear encoded, mitochondrial-targeted ATP6 and ATP8. * p<0.05; **p<0.01; ****p<0.0001; ns – not significant

Cytosolic and mitochondrial protons contribute to ETC function, which cooperatively builds and consumes a proton gradient to synthesize ATP. We therefore sought to evaluate how the ETC contributes to acidosis-driven MondoA activity. We used 143Bρ^0^ osteosarcoma cells, which lack mitochondrial DNA (mtDNA) and are respiration deficient (King and Attardi, 1989). TXNIP was induced in parental 143B cells treated with DMEM^Acidic^ (Figure 4A), yet the induction of TXNIP was blunted in 143Bρ^0^ cells (Figure 4B). TXNIP induction was rescued in 143Bρ^0^ cells that had been repopulated with wild type mitochondria (143Bρ^0^:WT-cybrid cells; Figure 4C). These genetic experiments confirm previous inhibitor studies that implicated a functional ETC in MondoA transcriptional activity (Yu et al., 2010, Han and Ayer, 2013).

Given the predominant role of the ETC in ATP synthesis, we determined whether mitochondrial ATP (mtATP) synthesis is required to trigger the MondoA/TXNIP axis. We used 143Bρ^0^:ΔATP6/ΔATP8 cybrid cells which have a point mutation in mtDNA that disrupts expression of both ATP6 and ATP8, required components of the F_0_F_1_-ATPase (ATP synthase, Figure 4D)(Boominathan et al., 2016, Jonckheere et al., 2008). Low pH-driven TXNIP expression was blunted in these cells, yet was partially rescued in cells with nuclear-encoded, mitochondrially-targeted ATP6 and ATP8 (Figure 4E)(Boominathan et al., 2016). These results indicate that mtATP synthesis is necessary for low pH to induce MondoA transcriptional activity. Consistent with this hypothesis, the ATP synthase inhibitor oligomycin completely blocked TXNIP induction in response to DMEM^Acidic^ (Figure 4 – figure supplement 1B).

### Acidosis drives the synthesis of mitochondrial ATP

ETC complexes I-IV build a proton gradient by pumping protons from the mitochondrial matrix to the inner membrane space. Given that the outer mitochondrial membrane is freely permeable to protons (Cooper, 2000), we hypothesized that acidosis leads to intracellular acidification, hyperpolarization of the inner-mitochondrial membrane and ATP synthesis. Using the pH-sensitive dye BCECF-AM, we determined that DMEM^Acidic^ treatment shifted intracellular pH from 7.2 to 6.5 (Figure 5A). The drop in pH was accompanied by an increase in mitochondrial membrane potential as measured by the dye JC1 (Figure 5B) and an increase in total cellular ATP levels (Figure 5C). Collectively these data show that treating cells with low pH medium increases total cellular ATP levels.

**Figure 5.**
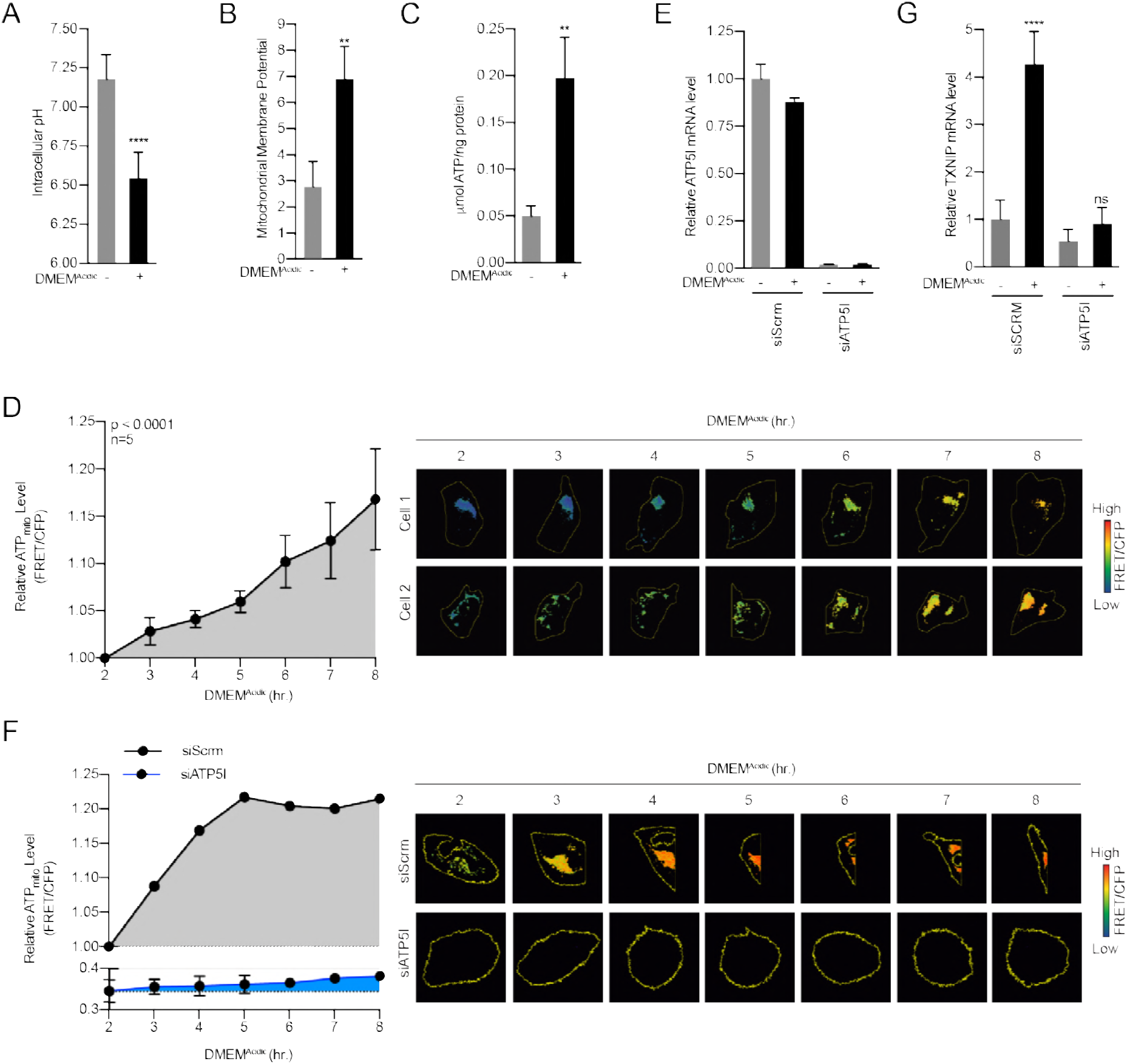
Acidosis drives the synthesis of mitochondrial ATP. (**A**) Intracellular pH of HeLa cells treated with DMEM^Acidic^ as determined by BCECF-AM staining. (**B**) Mitochondrial membrane potential was determined by JC1 staining. (**C**) Total cellular ATP levels were determined using luciferase-based assay. (**D**) Mit-ATEAM, a mitochondrial-targeted ATP-biosensor, was used to determine how DMEM^Acidic^ affects mitochondrial ATP. Widefield microscopy was used to capture images in the FRET and CFP channels. After images were obtained, mitochondria were analyzed for FRET and CFP signal. FRET signal was normalized using CFP. (**E**) ATP5I mRNA level in HeLa cells expressing scrambled (siSCRM, n=1) or ATP5I-specific siRNA (siATP5I, n=2). (**F**) Mit-ATEAM was used to determine how DMEM^Acidic^ affects mitochondrial ATP production in the context of siSCRM or siATP5I. (**G**) TXNIP mRNA level following DMEM^Acidic^ treatment of HeLa cells expressing scrambled or ATP5I-specific siRNA. ** p<0.01; ****p<0.0001; ns – not significant

Metabolite pools from whole cells can be vastly different from those observed in specific organelles (Abu-Remaileh et al., 2017, Chen et al., 2016). We therefore sought to determine how low pH affects mtATP levels. To accomplish this, we used a mitochondrial-targeted fluorescence resonance energy transfer (FRET) ATP biosensor (Mit-ATEAM, Figure 5 – figure supplement A-B). This biosensor consists of cp173-Venus fused to mseCFP via an ATP-binding linker region (Imamura et al., 2009). As a control, we used constructs with mutations in the ATP-binding linker that prevented ATP binding and FRET (Figure 5 – figure supplement 1C). HeLa cells treated with DMEM^Acidic^ showed increased FRET over time, indicating that acidosis drives an increase in mtATP but not cytosolic ATP (Figure 5D, Figure 5 – figure supplement 1C-E).

We next sought to determine whether the accumulation of mtATP resulted from increased synthesis or decreased mitochondrial export. We blunted expression of ATP5I, an essential component of the ATP synthase, using siRNA-mediated knockdown (Figure 5E). Consistent with our working model, ATP5I knockdown decreased not only the steady state level of mtATP, but also the low pH-driven increase in mtATP (Figure 5F). Furthermore, ATP5I knockdown prevented TXNIP induction in response to DMEM^Acidic^ treatment (Figure 5G). Together these data show that acidosis drives mtATP production through ATP synthase and that mtATP synthesis is required for low pH-driven MondoA transcriptional activity.

### MondoA senses G6P produced by mitochondrial-hexokinase

How might MondoA sense mtATP? MondoA, Mlx and hexokinase 2 (HK2) are all resident at the outer mitochondrial membrane (Figure 6A-B) (Robey and Hay, 2006, Sans et al., 2006). Mitochondria-bound HK2 has preferential access to mtATP that is exported from the mitochondria (Wilson, 2003). The enzymatic activity of HK2 transfers the terminal phosphate from ATP to glucose to generate G6P. Because G6P is a known activator of MondoA transcriptional activity, we speculated that acidosis-induced mtATP drives the synthesis of G6P to trigger MondoA transcriptional activity (Figure 6B). We have tested this model in several ways. First, we determined how acidosis alters steady-state metabolite levels. Consistent with reports showing that acidosis leads to increased mitochondrial metabolism (Lamonte et al., 2013, Chen et al., 2008, Dietl et al., 2010), DMEM^Acidic^ drove an increase in TCA cycle intermediates (Figure 6C). By contrast, most glycolytic intermediates were decreased in response to DMEM^Acidic^; however, G6P levels were increased 3-fold (Figure 6C).

**Figure 6.**
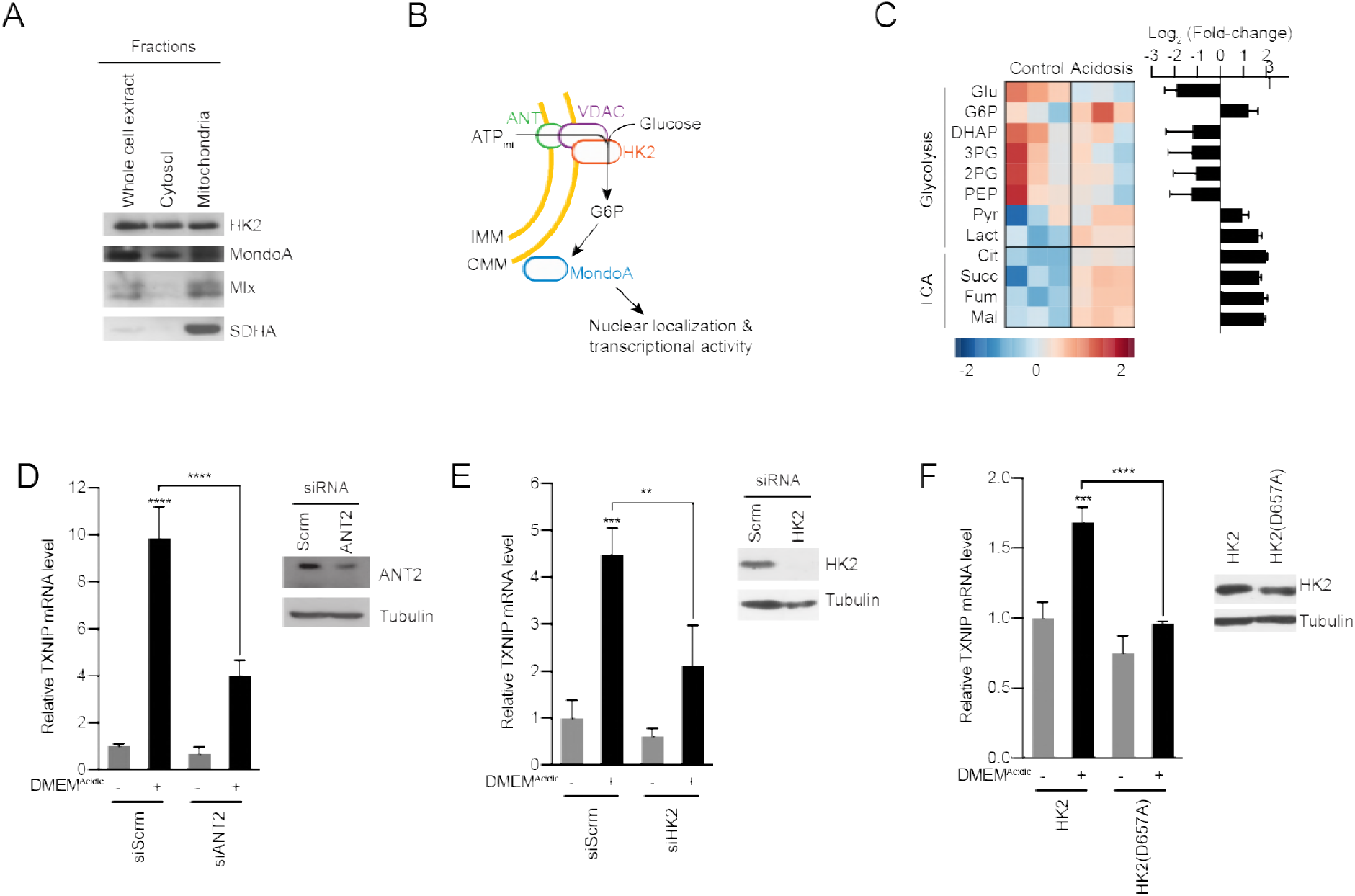
MondoA senses G6P produced by hexokinase utilization of mtATP. (**A**) Cellular fractionation of BJ-Tert cells indicating mitochondrial localization of HK2, MondoA and Mlx. Succinate dehydrogenase A (SDHA) serves as a control for the mitochondria fraction. (**B**) Schematic illustrating how mtATP could contribute to MondoA transcriptional activity. As mtATP is exported from the mitochondria, it is used as a substrate to produce G6P by mitochondrial-bound HK2, resulting in MondoA activation. (**C**) Heatmap and log_2_ fold-changes of glycolytic and TCA metabolites measured using GC-MS. TXNIP mRNA levels of HeLa cells treated with DMEM^Acidic^ and expressing a pool of four siRNAs against (**D**) ANT2 and (**E**) HK2, or (**F**) expressing HK2 and HK2(D657A). **p<0.01; ***p<0.001; ****p<0.0001; ns – not significant

Second, we tested the contribution of the channel, comprised of the adenine-nucleotide transporter (ANT) in the inner-mitochondrial membrane and voltage-dependent anion channel (VDAC) in the outer-mitochondrial membrane, that exports mtATP from the mitochondria. Consistent with our working model, which states that mtATP must be exported from the matrix, siRNA-mediated knockdown of ANT2 prevented TXNIP induction in response to low pH medium (Figure 6D). This finding suggests that mtATP functions outside the mitochondria to trigger MondoA transcriptional activity, rather than by an indirect signaling-based mechanism.

Third, we used several approaches to test the contribution of HK2 to low pH-driven MondoA activity. siRNA pools against HK2 blocked TXNIP induction in response to low pH treatment (Figure 6E), demonstrating a requirement for HK2. Further, overexpression of HK2(D657A), which lacks kinase activity (Arora et al., 1991), blocked the induction of TXNIP in response to low pH (Figure 6F), supporting the notion that synthesis of G6P is critical for the induction of MondoA transcriptional activity.

Fourth, we tested the contribution of mitochondria-localized HK2 to low pH-driven MondoA transcriptional activity. HK2 localizes to the outer-mitochondrial membrane via interactions with VDAC (Wilson, 2003). Ectopic expression of mVDAC(E72Q), a mutant mouse orthologue of VDAC1, prevents the interaction between VDAC and HK2 (Abu-Hamad et al., 2008, Zaid et al., 2005), blocked the mitochondrial localization of HK2 as expected and completely blocked TXNIP induction in response to DMEM^Acidic^ (Figure 7A). We complemented these loss-of-function experiments with a gain-of-function approach designed to determine whether mitochondrial localization of hexokinase was sufficient for low pH-induced MondoA transcriptional activity. To accomplish this goal, we artificially tethered HK2 to the mitochondria. We achieved this by fusing VDAC1 to the first 10 β-strands of GFP (mVDAC1-GFP(1-10)) and by fusing HK2 to the last β-strand of GFP (HK2-GFP(11)). When co-expressed, the β-strands of GFP self-assemble (Kamiyama et al., 2016), linking mVDAC1 and HK2 (Figure 7B). Expression of mVDAC1(E72Q)-GFP(1-10), which does not interact with HK2, blocked TXNIP induction (Figure 7B-C). However, co-expression of mVDAC1(E72Q)-GFP(1-10) and HK2-GFP(11) rescued HK2 mitochondrial localization and TXNIP induction (Figure 7B-C). Together these data show that mitochondrial-localized HK2 is both necessary and sufficient for acidosis-driven MondoA activity.

**Figure 7.**
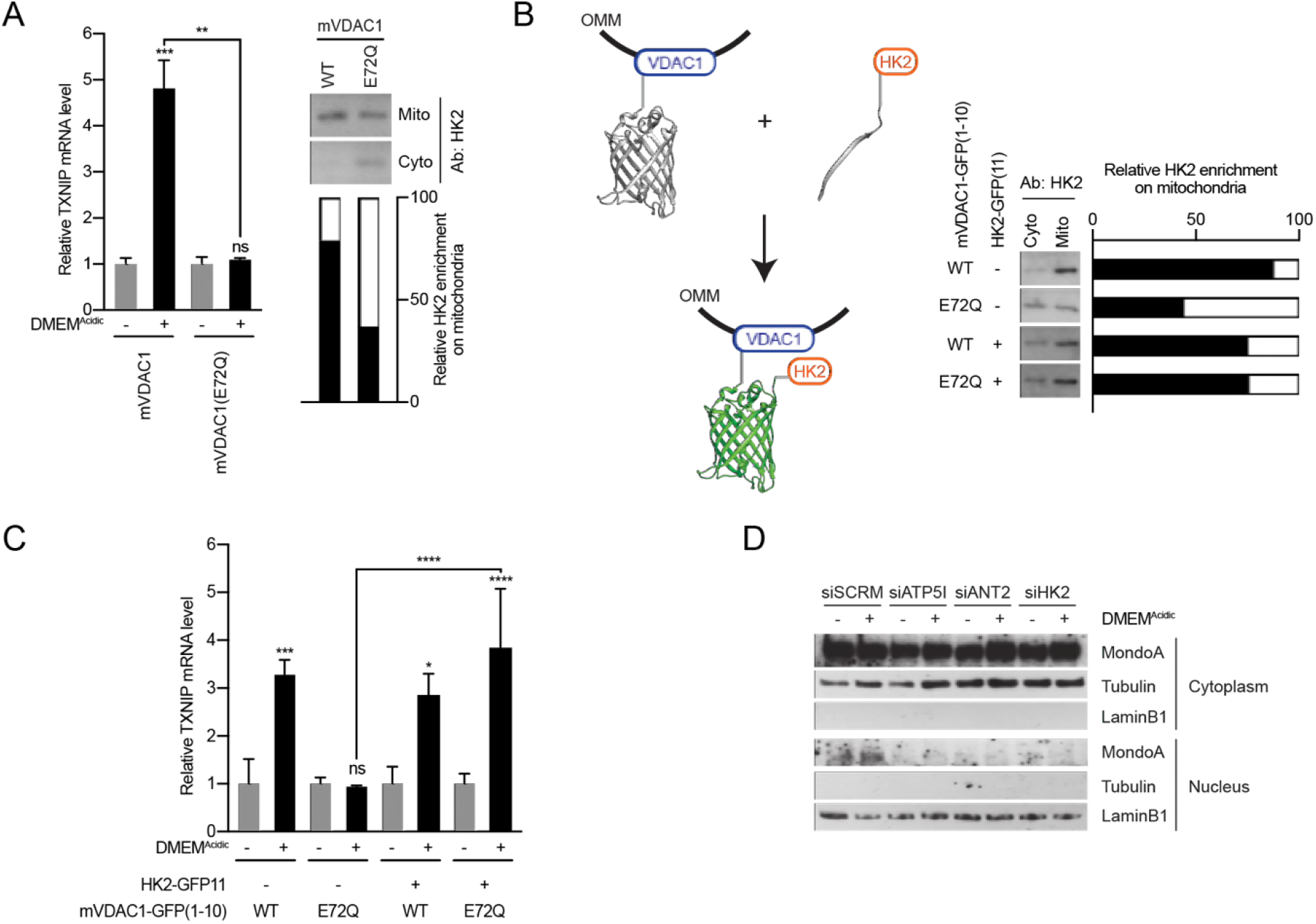
MondoA senses G6P produced by mitochondrial-bound hexokinase. (**A**) TXNIP mRNA levels in BJ-Tert cells expressing mVDAC1-GFP and mVDAC1(E72Q)-GFP and treated with DMEM^Acidic^ HK2 localization was also analyzed by cellular fractionation and densitometry was used to quantify the relative amount of HK2 on the mitochondria. Of note, HK2 became increasingly enriched in the cytoplasmic fraction. (**B**) Schematic depicting the use of GFP(1-10) and GFP(11) to artificially tether mVDAC1 and HK2. HK2 localization was also analyzed by cellular fractionation and densitometry was used to quantify the relative amount of HK2 on the mitochondria. (**C**) TXNIP mRNA levels of BJ-Tert cells treated with DMEM^Acidic^ and expressing mVDAC1-GFP, mVDAC1(E72Q)-GFP and HK2-GFP(11). (**D**) MondoA nuclear localization was determined by cellular fractionation of HeLa cells treated with DMEM^Acidic^ and with siSCRM (siRNA control), siATP5I, siANT2 and siHK2. Tubulin and LaminB1 served as controls for cytoplasm and nuclei, respectively. * p<0.05; **p<0.01; ***p<0.001; ****p<0.0001; ns – not significant

Finally, we determined the effects of acidosis-driven mtATP and G6P synthesis on MondoA nuclear localization. While the majority of MondoA resides in the cytosol (Figure 7D), DMEM^Acidic^ treatment drove an increase in MondoA nuclear localization in cells treated with an siRNA control (Figure 7D); however, MondoA nuclear accumulation was blunted in cells treated with siRNA pools against ATP5I, ANT2 and HK2 (Figure 7D). These finding demonstrate that mtATP and G6P synthesis are required for MondoA to accumulate in the nucleus in response to low pH (Figure 8).

**Figure 8.**
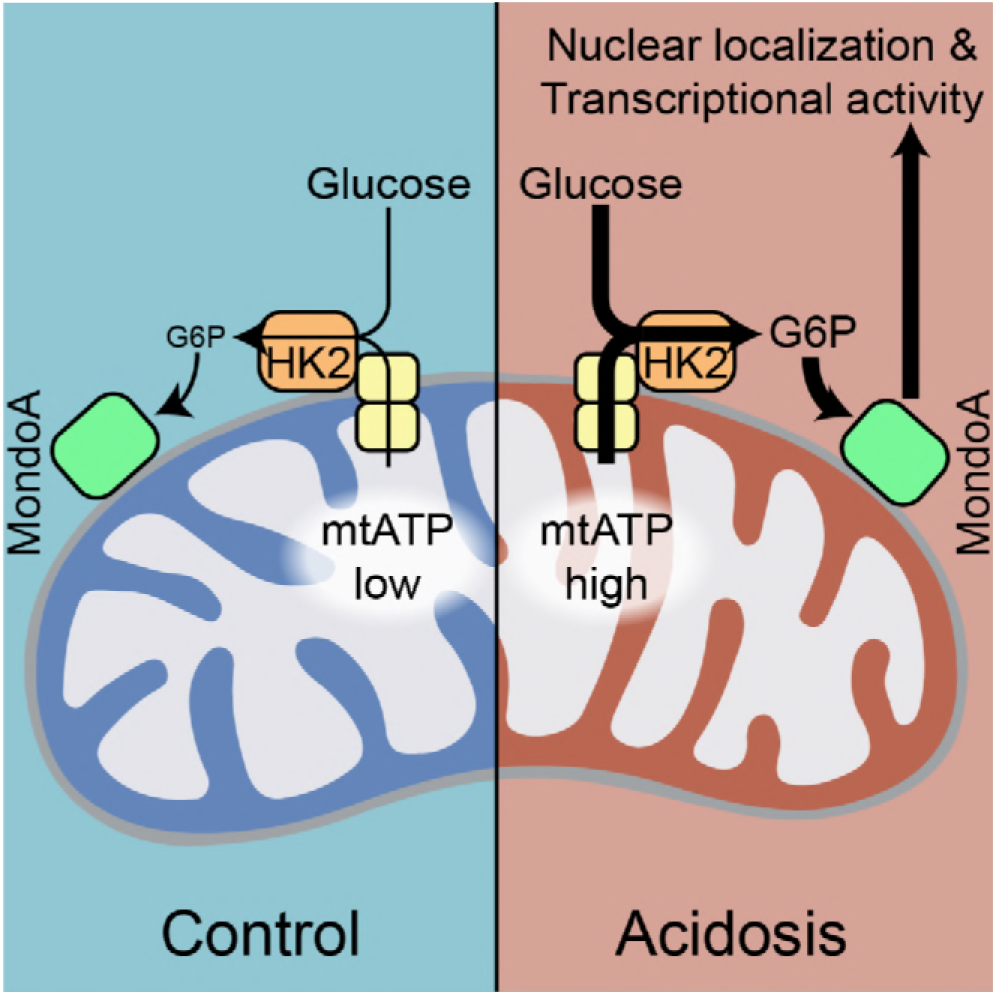
Model. Schematic depicting how acidosis drives MondoA transcriptional activity through the generation of mitochondrial ATP and utilization by mitochondria-bound hexokinase to produce G6P, which drives MondoA nuclear localization and transcriptional activity.

## DISCUSSION

Previous studies established that MondoA’s transcriptional activity is highly-dependent on two signals: glucose and a signal from the ETC. By dissecting how low pH drives MondoA transcriptional activity, we establish here that the ETC signal is mtATP. Previous studies showed that a functional ETC is required for basal TXNIP expression, thus we propose that mtATP is a general requirement for MondoA transcriptional activity. Via the activity of OMM-bound HK2, mtATP couples to cytoplasmic glucose to generate G6P, which drives the nuclear accumulation and transcriptional activity of MondoA:Mlx complexes (Figure 8). Further, we previously demonstrated (Sans et al., 2006), and confirmed here (Figure 4A), that MondoA and Mlx also interact with the OMM. Therefore, we propose that MondoA:Mlx and HK2 constitute a sensing and response module that integrates signals from the cytoplasm and the mitochondria to coordinate the transcriptional response to the cells two predominant energy sources. We studied how acidosis drives mtATP production and MondoA transcriptional activity. It will be interesting to determine whether other cellular signals that drive MondoA transcriptional activity also function by controlling mtATP pools.

By binding the mitochondria, hexokinase has increased specific activity and decreased feedback inhibition by G6P (Robey and Hay, 2006). By localizing to the mitochondria and sensing G6P derived from mitochondria-bound hexokinase, we propose that MondoA activity is coupled to mitochondrial hexokinase activity and mtATP synthesis. Given the OMM localization of MondoA, Mlx and HK2, we suggest that the OMM serves as a scaffold for nutrient sensing by MondoA, akin to other nutrient sensors that are tethered to organellar membranes, e.g. the mTORC1 complex is tethered to the lysosome where it integrates intra-lysosomal nutrient levels and cytosolic growth factor signals to control biosynthesis (Wolfson and Sabatini, 2017), and the SREBP/Scap complex which is resident in the ER membrane where it monitors cholesterol and oxysterol availability and controls a transcriptional response to low sterol levels (Moon, 2017).

Our data suggest that MondoA functions as a coincidence detector which simultaneously senses mtATP and glucose through the synthesis of G6P (Figure 8). Such a model ensures that the availability of glucose is tightly linked to mitochondrial activity and ATP synthesis. Conceptually, by coupling mtATP and cytosolic glucose, MondoA functions as a sensor of high cellular energy charge and via its transcriptional regulation of TXNIP, and potentially other targets, restricts glucose uptake and aerobic glycolysis to restore energy balance. Further, high TXNIP levels are correlated with oxidation of triglycerides, branched chain amino acids and lactate (DeBalsi et al., 2014, Bodnar et al., 2002), suggesting an additional role for the MondoA/TXNIP axis in driving utilization of non-glucose fuels when cellular energy charge is high. The precise mechanistic details of how MondoA senses G6P and the impact on cell metabolism remains to be clarified; however, the current data is most consistent with a direct allosteric model where G6P binds MondoA directly (McFerrin and Atchley, 2012, Peterson et al., 2010, Li et al., 2010).

Changes in intracellular pH have dramatic effects on cell function: generally acidic pH is anti-proliferative whereas alkaline pH is pro-proliferative. Acidic intercellular pH is correlated with an inhibition of aerobic glycolysis and a blockage of glucose uptake (Webb et al., 2011). Our data suggest that TXNIP induction contributes to this suppression of glucose metabolism driven by acidic pH. Further, an acidosis-dependent gene signature, of which TXNIP is a member correlates with better clinical outcomes in breast cancer (Chen et al., 2010, Chen et al., 2008). Interestingly, Otto Warburg noted that increased sodium bicarbonate and alkaline pH favor glycolysis (Koppenol et al., 2011, Warburg, 1925). Intracellular alkalization is now a widely accepted hallmark of cancer metabolism (Webb et al., 2011), that has pleiotropic effects on tumorigenesis, the most predominant being a transition from oxidative metabolism to aerobic glycolysis (Reshkin et al., 2000). Because, MondoA transcriptional activity at the TXNIP promoter is suppressed by high sodium bicarbonate and alkaline pH (Figure 2 – figure supplement 1C), we propose that TXNIP down regulation contributes to the shift to high glucose uptake driven by alkaline pH. It will be important to determine whether alkaline pH suppresses mitochondrial function and restrict mtATP production. Collectively, our data suggest that MondoA/TXNIP axis plays a critical role in how cancer cells sense and respond to dysregulated pH.

Our gene expression data demonstrates that MondoA is essential for the regulation of 37% of an acidosis-driven transcriptional response. Among the MondoA-dependent genes are fatty acid and mitochondrial metabolism genes. Given that these pathways are enhanced by acidosis (Corbet et al., 2016, Lamonte et al., 2013, Khacho et al., 2014), we propose that MondoA plays a critical role in an adaptive metabolic response to acidosis. Most genes were only partially dependent on MondoA; however, TXNIP and ARRDC4 were entirely dependent on MondoA, suggesting that TXNIP and ARRDC4 are direct MondoA targets whereas the other targets may be regulated by indirect mechanisms. Consistent with this finding, MondoA is enriched on the promoters of TXNIP and ARRDC4 in MDA-MB-231 cells, but not on the promoters of the other acidosis-regulated genes identified here (data not shown).

Finally, it is well established that oncogenes drive a shift from oxidative metabolism to aerobic glycolysis (Pavlova and Thompson, 2016). The resulting shift away from ATP synthesis in the mitochondria to ATP synthesized by glycolysis in the cytosol would be predicted to restrict MondoA-dependent activation of TXNIP expression and reinforce glucose uptake and aerobic glycolysis. Strikingly, TXNIP expression is also downregulated by a variety of pro-growth signals such as mTOR, PI3K, Ras and Myc (Elgort et al., 2010, Kaadige et al., 2015, Shen et al., 2015), which results in increased glucose uptake. Together these two findings place MondoA and it regulation of TXNIP both upstream and downstream of metabolic reprogramming towards aerobic glycolysis.

## MATERIALS AND METHODS

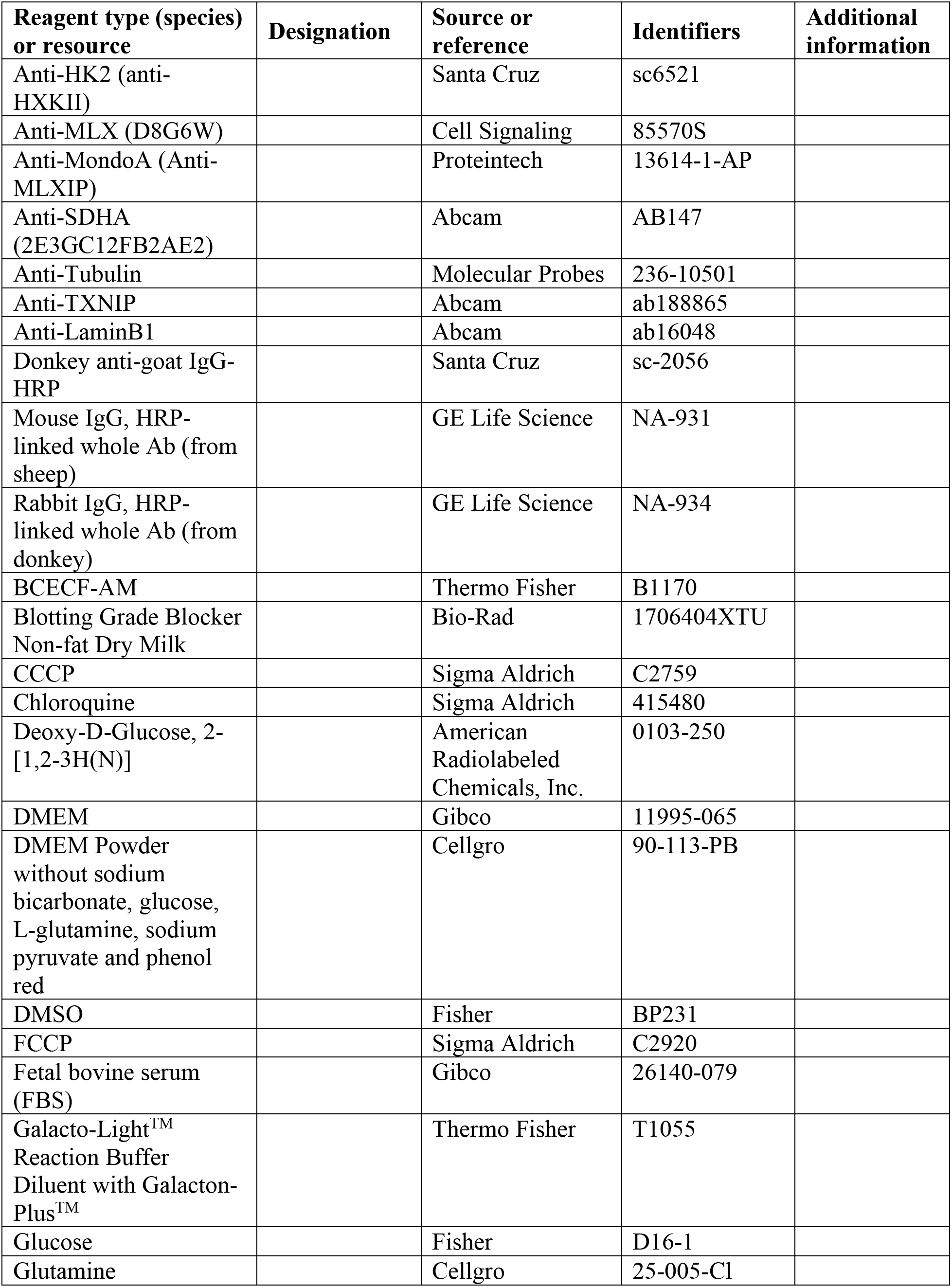
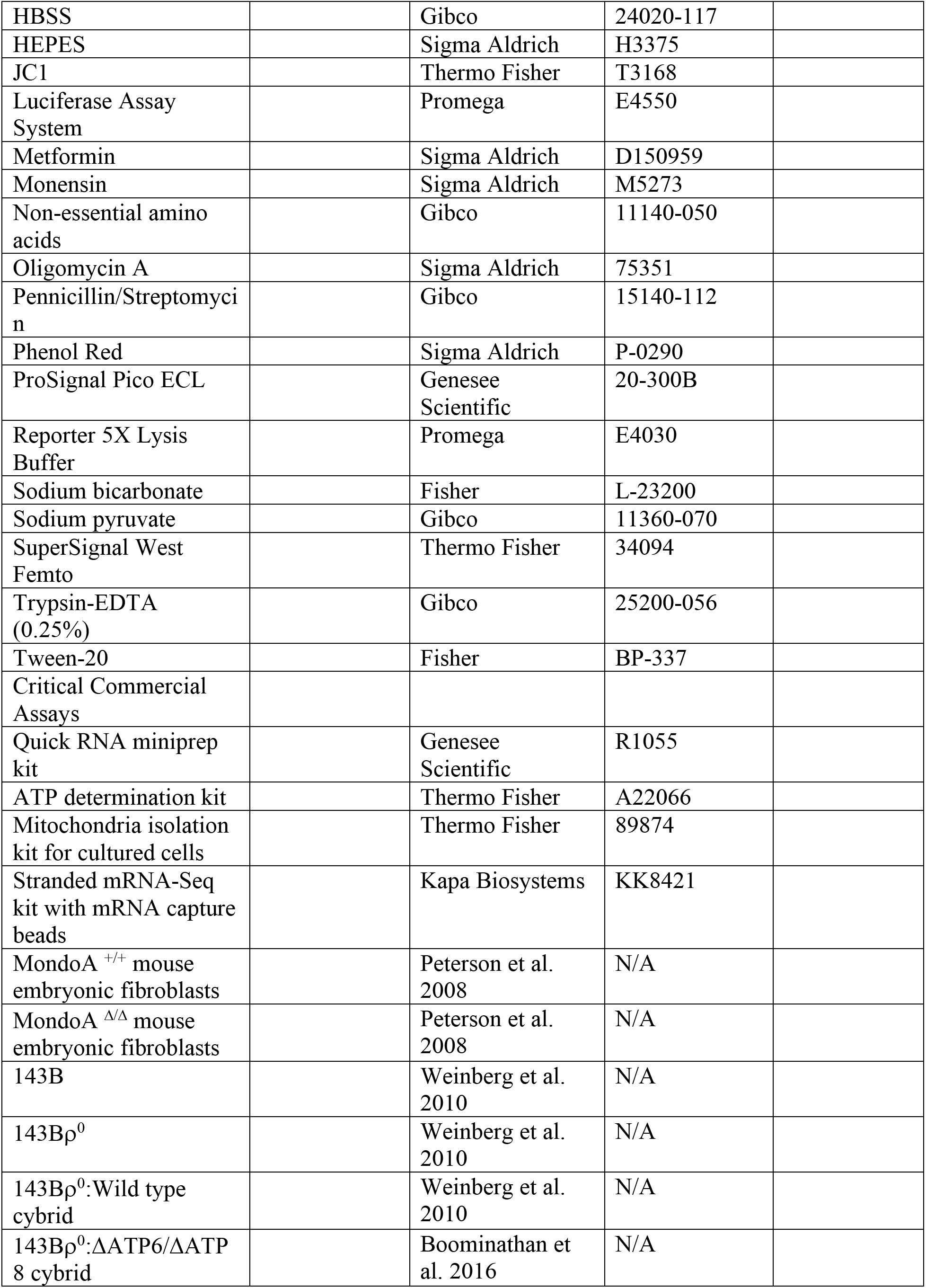
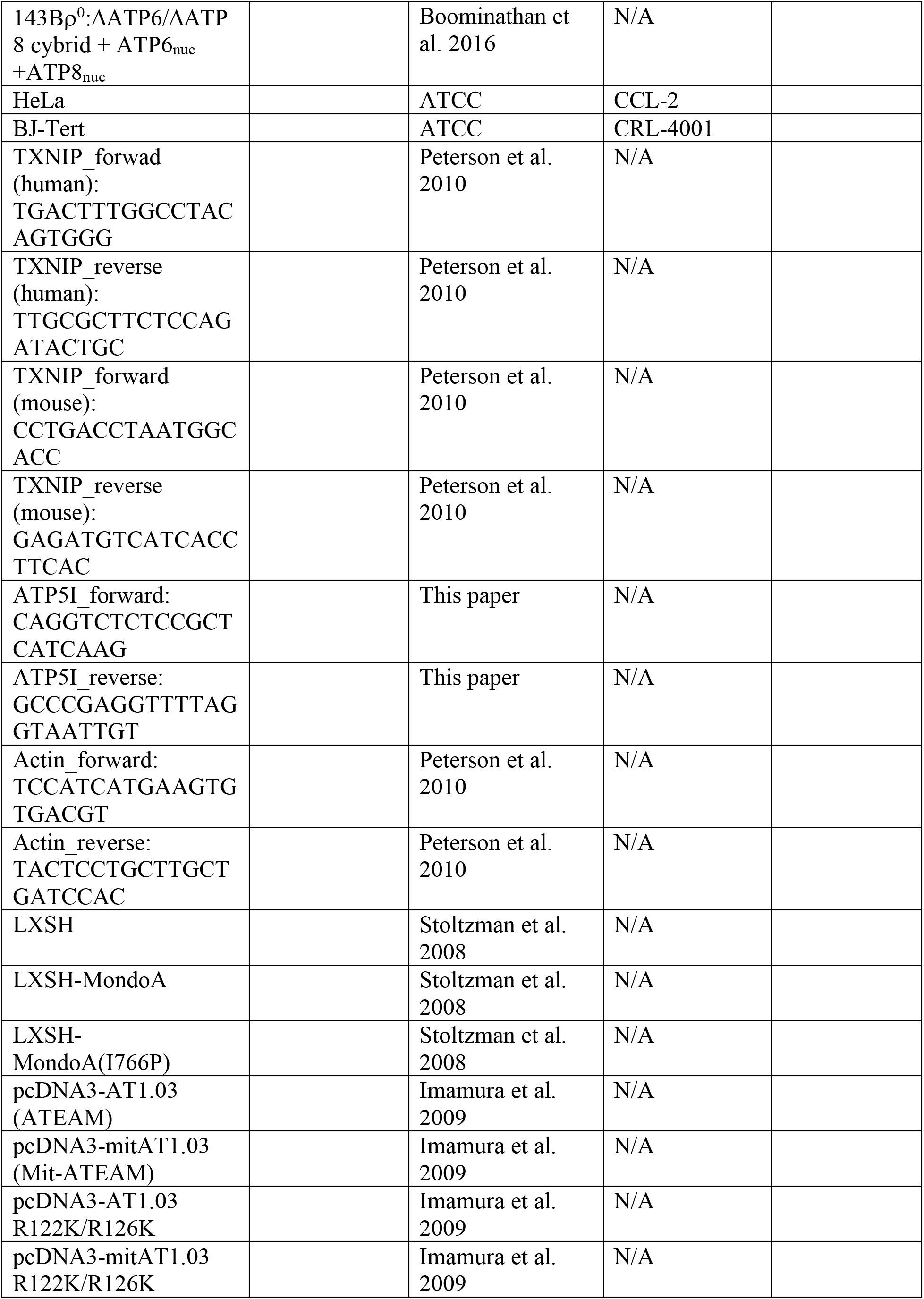
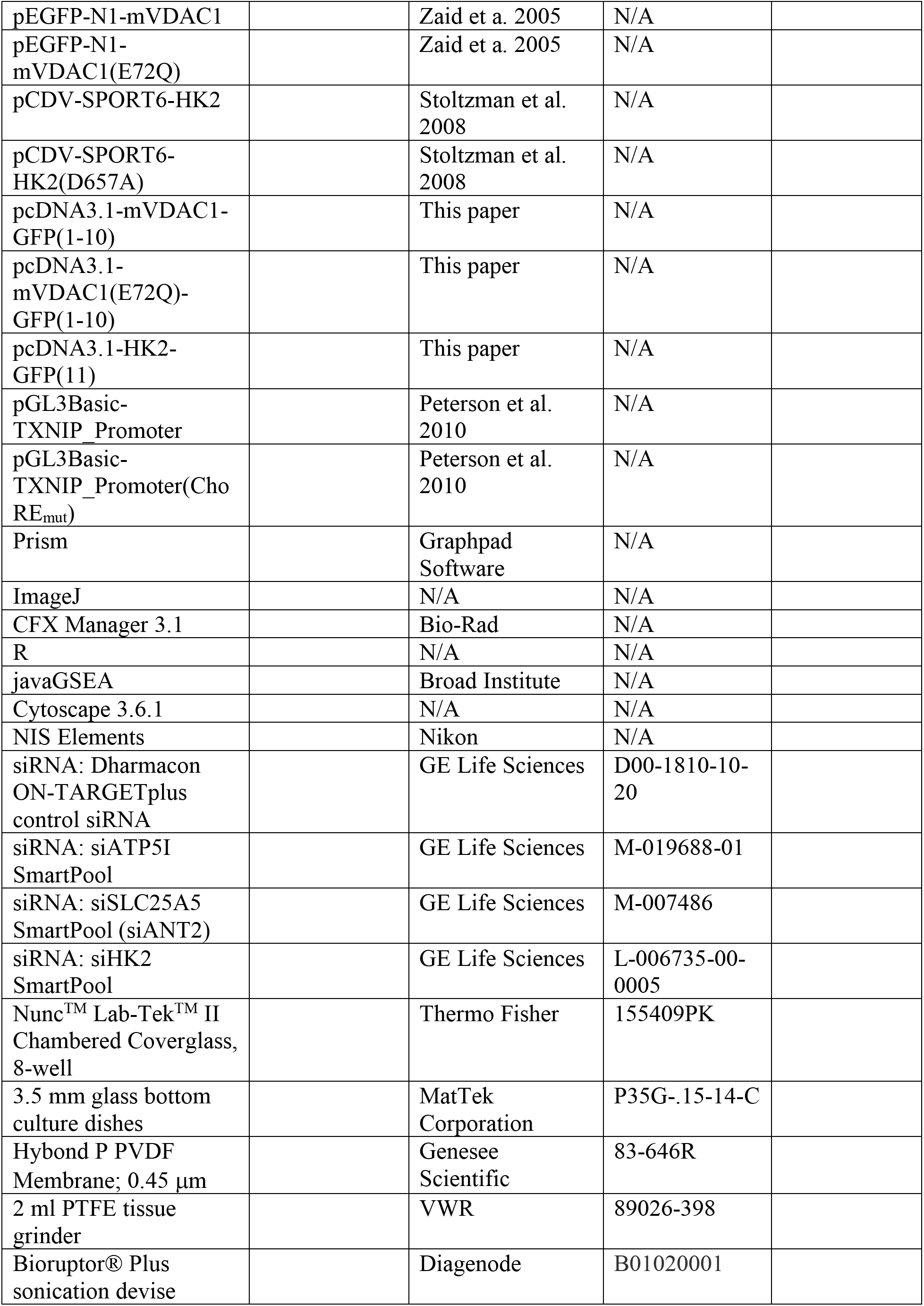
Key Resources Table

### Cell lines

A list of cell lines used is provided in the Key Resources Table. All cells were maintained in DMEM +10% FBS (Gibco), 100 units/mL penicillin (Gibco) and 100 units/mL streptomycin (Gibco). 143Bρ^0^ and cybrids were cultured with 1 mM sodium pyruvate and 50 μg/mL uridine. Cells were passaged and treated in an incubator set at 37 °C and 5% CO_2_.

HeLa:MondoA-KO cells were generated by expressing CRISPR/Cas9, three sgRNAs (GeCKO library 2.0) and a homology-directed repair (HDR) construct containing a puromycin-resistance cassette (Santa Cruz Biotechnology). HDR incorporation into the genome was determined by selecting for cells resistant to 2.5 μg/mL puromycin. Loss of MondoA was determined by immunoblotting.

### Treatments

HBSS was supplemented with glucose to 20 mM prior to treatment. Low pH treatment media (pH 6.5) was prepared from DMEM powder without glutamine, glucose, pyruvate, sodium bicarbonate and phenol red (DMEM^Acidic^). The following were added: glutamine to 2 mM, glucose to 20 mM, pyruvate 1 mM, sodium bicarbonate to 0.35 g/L and phenol red to 16 mg/L. For live cell imaging, phenol red was omitted.

### Plasmid construction

Plasmids were created using either standard restriction digest and ligation or Gibson assembly (NEB). A list of plasmids used, the vector backbone and their source is provided in the Key Resources Table.

### Quantitative PCR

Total cellular RNA was extracted using a Quick RNA Miniprep Kit (Zymo Research) according to manufacturer’s recommendations. cDNA was synthesized from 200 ng mRNA using the GoScript Reverse Transcription System (Promega) with oligo-dT primers. A 100-fold dilution was used in a PCR reaction containing SYBR Green and analyzed on a CFX Connect Real Time System. Values were determined using a standard curve. For each sample, three technical replicates were performed and averages determined.

### Immunoblotting

Equal concentrations of denatured protein lysates were resolved on 10% SDS-PAGE gel with a stacking gel. Proteins were electrotransferred to PDVF membrane (Genesee Scientific). Membranes were incubated in 5% (weight/volume) blotting-grade non-fat dry milk (Bio-Rad) in TBST (Tris-buffered saline, pH 7.4 and 0.1% Tween-20) for 30 minutes at room temperature with gentle rocking. Membranes were then transferred to antibody-dilution buffer (20 mM Tris, pH 8.0; 200 mM NaCl; 0.25% Tween-20; 2% bovine serum albumin; 0.1% sodium azide) and incubated for one hour at room temperature or overnight at 4 °C with gentle rocking. Membranes were washed with TBST and vigorous rocking at room temperature. Membranes were then incubated in secondary antibody diluted in 5% (weight/volume) blotting-grade non-fat dry milk (Bio-Rad) in TBST for one hour at room temperature with gentle rocking. Membranes were then washed again and proteins were detected with chemiluminescence using standard or high sensitivity ECL (Genesee Scientific or Thermo Fisher, respectively). Antibodies were used at the following dilutions: Anti-GFP 1:1000; Anti-HK2 1:1,000; Anti-Mlx 1:1,000; Anti-MondoA 1:2,000; Anti-SDHA 1:15,000; Anti-Tubulin 1:50,000; Anti-TXNIP 1:2,000; Anti-goat HRP 1:20,000; Antimouse HRP 1:5,000 and Anti-rabbit HRP 1:15,000.

### ATP quantification

After treatment, cells were washed once with cold PBS. Cells were scraped into boiling TE buffer (1 mL per 3.5 cm dish), which was collected into 1.5 mL centrifuge tube. Cells were then boiled for 5 min. Lysates were cleared by centrifugation at 20,000xg for 5 minutes. The ATP determination kit (Thermo Fisher) was used with 10 μL of supernatant. A standard curve was generated using purified ATP.

### Live cell imaging (mtATP determination): Widefield microscopy

Widefield microscopy was used for Figure 5 and Figure 5 – figure supplement 1. Cells were plated on 3.5 mm glass bottom culture dishes (MatTek Corporation). The following day 100 ng Mit-ATEAM was transfected using Lipofectamine 3000 (Thermo Fisher) according to manufacturer’s recommendations. The next day cells were treated with DMEM^Acidic^ lacking phenol red. Real time live imaging was conducted for 8 hours using a Nikon A1R with a 40X lens. For each time point images were captured using 488/525 (YFP), 405/480 (CFP), and 405/525 (FRET) excitement/emission (nm).

Images were analyzed using ImageJ. We used the YFP channel to identify and isolate mitochondrial regions for each image. We isolated these same regions from the CFP and FRET channel. Total intensity was determined for each image. FRET/CFP ratios were determined and normalized to the 2-hour time point. RatioPlus was used to make pseudo-colored images.

### Live cell imaging (mtATP determination): Confocal microscopy

Confocal microscopy was used for Figure 5 – figure supplement 1. Cells were plated on 8-well Nunc™ Lab-Tek™ II Chambered Coverglass (Thermo Fisher). The following day 200 ng DNA was transfected using Lipofectamine 3000 (Thermo Fisher) according to manufacturer’s recommendations. The next day cells were treated DMEM^Acidic^ lacking phenol red or DMEM lacking phenol red with CCCP. Real time live imaging was conducted for 8 hours using a Nikon A1 with a 20X lens. For each time point images were captured using 402/488 (CFP) and 402/525 (FRET) excitement/emission (nm). Images were analyzed using ImageJ. RatioPlus was used to make pseudo-colored images. Total intensity for each image was determined.

### Glucose uptake

Cells were incubated with deoxy-D-glucose-2[1,2-3H(N)] (American Radiolabeled Chemicals, Inc.) in Krebs-Ringer-HEPES buffer (NaCl,116 mM; KCl, 4 mM; MgCl_2_, 1 mM; CaCl_2_, 1.8 mM; 2-deoxy-D-glucose, 20 mM; HEPES pH 7.4, 10 mM) for 10 minutes. Cells were then washed, harvested and analyzed for radioactivity using a scintillation counter. A standard was used to determine the exact molar content in each sample. deoxy-D-glucose-2[1,2-3H(N)] was normalized to protein content as determined by a Bradford Protein Assay (Bio-Rad).

### Mitochondrial membrane potential

Cells were plated on 8-well Nunc™ Lab-Tek™ II Chambered Coverglass (Thermo Fisher). 45 minutes prior to experiment, cells were loaded with JC1 (1 μg/mL). Cells were treated DMEM^Acidic^ lacking phenol red or DMEM lacking phenol red. A Nikon A1 confocal and NIS Elements AR were used to capture images. For each time point images were captured by exciting with 488 nm light and reading the emission at 530 nm (green) and 595 nm (red). The ratio of red to green was used to quantify changes in membrane potential one hour after acidosis treatment.

### Intracellular pH

Cells were plated on 3.5 mm glass bottom culture dishes (MatTek Corporation). The next day cells were treated with normal or low pH DMEM for 4 hours. Cells were treated with BCECF-AM (1 μM) 30 minutes prior to the end of the experiment. A standard curve was generated by treating cells with media of varying pHs and Nigericin (5 μM), which equilibrates intracellular and extracellular pH. A Nikon A1 confocal and NIS Elements AR were used to capture images by exciting with 488 nm and reading emission at 530 nm and 595 nm. The 595/530 nm fluorescence emission ratio was used to generate a calibration curve and determine intracellular pH for acidosis-treated cells.

### Mitochondria purification

Mitochondria were purified from ~20×10^6^ cells using a Mitochondria Isolation Kit for Cultured Cells (Thermo Fisher). Cells were processed using a PTFE tissue grinder (VWR). Following purification, mitochondria were resuspended in 100 μl radioimmunoprecipitation (RIPA) buffer. 100 μl of both mitochondria and cytosolic fractions were sonicated at using a Bioruptor sonication device (Diagenode). Sonication was performed 4°C using 30 second on/off pulses at the high setting. Following sonication, lysates were centrifuged and supernatants were collected and analyzed for protein content using a Bradford Protein Assay (Bio-Rad). 1-5 μg of sample were used for immunoblot analysis.

### Subcellular Fractionation: Nuclei and cytoplasm

Three days prior to fractionation siRNAs were transfected using Lipofectamine 3000 (Thermo Fischer). Cells were washed with cold PBS and dislodged from plate by scraping. Cells were pelleted by centrifugation and resuspended in 1 mL of fractionation buffer (40 mM HEPES pH 7.9, 137 mM NaCl, 2.7 mM KCl, 1.5 mM MgCl_2_, 0.34 M sucrose, 10% glycerol, 1 mM DTT, 0.5% NP40, protease and phosphatase inhibitors). Cells were incubated on ice for 10 minutes then pelleted by centrifugation at 1000 rcf for 5 minutes. The supernatant was kept (cytoplasm) and the pellet (nuclei) was washed three times with 0.5 mL fractionation buffer.

### Luciferase Assay

Cells were seeded and the next day transfected with constructs containing a 1518-bp fragment of the TXNIP promoter (or a mutant)-driven luciferase and CMV-driven beta-galactosidase (Kaadige et al., 2009). Cells were harvested in 1X Buffer RLB (Promega). Luciferase was detected using the Luciferase Detection System (Promega) and beta-galactosidase was detected using Galacto-Light™ Reaction Buffer Diluent with Galacto-Plus™ Substrate (Thermo Fisher). Luminescence was determined using a GloMax 96 Microplate Luminometer (Promega). Luciferase values were normalized to beta-galactosidase.

### GC-MS

Following treatment, cells were collected into a 1.5 mL microcentrifuge tube then snap frozen using liquid nitrogen. Cells were kept at −80°C until metabolite extraction was performed. 450 μL of cold 90% methanol and internal standards were added to cells and incubated at −20°C for 1 hour. Tubes were then centrifuged at −20,000×g for 5 minutes at 4°C. Supernatants were dried using a speed-vac.

Samples were converted into volatile derivatives amenable to GC-MS. Briefly, dried samples were resuspended in O-methoxylamine hydrochloride (40 mg/mL) then mixed with 40 μL N-methyl-N-trimethylsilyltrifluoracetamide and mixed at 37°C. After incubation, 3 μL fatty acid methyl ester standard solution was added. 1 μL of this final solution was injected into gas chromatograph with an inlet temperature of 250°C. A 10:1 split ratio was used. Three temperatures were ramped with a final temperature of 350°C and a final 3-minute incubation. A 30 m Phenomex ZB5-5 MSi column was used. Helium was used as carrier gas at 1 mL/minute. Samples were analyzed again with a 10-fold dilution.

Data was collected using MassLynx 4.1 software (Waters). Metabolites were identified and peak area was determined using QuanLynx. Data was normalized using Metaboanalyst 3.6 (http://www.metaboanalyst.ca/). Quantile normalization, log transformation and Pareto scaling were used. Normal distribution of values was used to determine fold changes.

### RNA-sequencing library construction and analysis

Total RNA was extracted from cells using a Quick RNA Miniprep Kit (Zymo Research) according to manufacturer’s recommendations. mRNA was isolated and library production performed using a Stranded mRNA-Seq Kit with mRNA Capture Beads (Kapa). Library quality was analyzed using an Agilent High Sensitivity D1000 ScreenTape. Single-end sequencing for 50 cycles was performed using an Illumina HiSeq. The resulting FASTQ files were aligned to the human genome (hg38) using STAR. DESeq2 was used to quantify transcript abundance, differential expression, FPKM values, and interaction terms (genotype:treatment combinatorial statistic).

Overrepresentation analysis performed using ConsensusPathDB. Pathway-based sets were analyzed from Wikipathways. A p-value cutoff of 0.01 and a minimum overlap of 2 genes was used. Enriched pathways were verified by comparing fold-changes obtained from DESeq2.

Gene set enrichment analysis and leading edge analysis (Broad Institute) was conducted using FPKM values and all gene sets from in the Molecular Signature Database. Leading edge analysis was visualized using the Cytoscape (p-value ≤ 0.001 and overlap coefficient ≥ 0.5).

### Gene signature

mRNA expression z-scores were obtained for 2509 breast cancer tumors (Pereira et al., 2016). Acidosis regulated genes were determined from the gene set GO_RESPONSE_TO_ACIDIC_PH in the Molecular Signature Database. Principal component analysis was conducted for all tumors using the expression levels of acidosis regulated genes. Gene signature scores were determined as the first principle component. This was compared to TXNIP expression for the same tumors.

The normalized expression (log_2_(normalized-counts + 1)) of TXNIP, SLC16A3 (MCT4), SLC16A1 (MCT1) and SLC9A1 (NHE1) was determined using the UCSC Xena browser. Spearman and Pearson coefficients were used to correlate gene expression. The following datasets were used: TCGA-BRCA, TCGA-LUNG, TCGA-GBM, GTEx-muscle and GTEx-skin.

### Quantification and statistical analysis

Data is presented as mean ± standard deviation. One-way ANOVA was used to account for variation and significance was determined using a two-tailed Student’s t-test. Unless otherwise indicated, at least three biological replicates were used for each analysis.

## ACKNOWLEDGEMENTS

We thank members of the Ayer Lab and Mahesh B. Chandrasekharan for helpful discussions and comments on this manuscript. We also thank Hiroyuki Noji (University of Tokyo), Varda Shoshan-Barmatz (Ben-Gurion University of Negev), Matthew S. O’Connor (SENS Research Foundation Research Center), and Jared Rutter (University of Utah) for reagents and advice. D.E.A. was supported by National Institutes of Health Grants 5RO1GM055668 and 1R01CA222650-01A1, by developmental funds from the Huntsman Cancer Foundation, and by Department of Defense Grant W81XWH1410445.

---

**Figure 1 – figure supplement 1.**
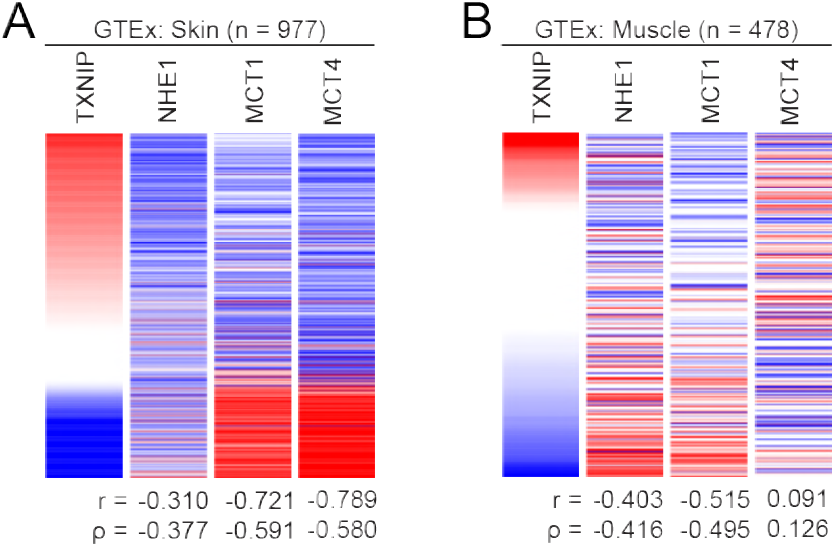
TXNIP correlates with genes that regulate intracellular pH. Heatmaps depicting the expression of TXNIP mRNA compared to MCT4, MCT1 and NHE1 for normal (**A**) skin and (**B**) muscle tissues. All expression data was collected from GTEx. Spearman and Pearson correlation statistics are reported as r and ρ, respectively.

**Figure 2 – figure supplement 1.**
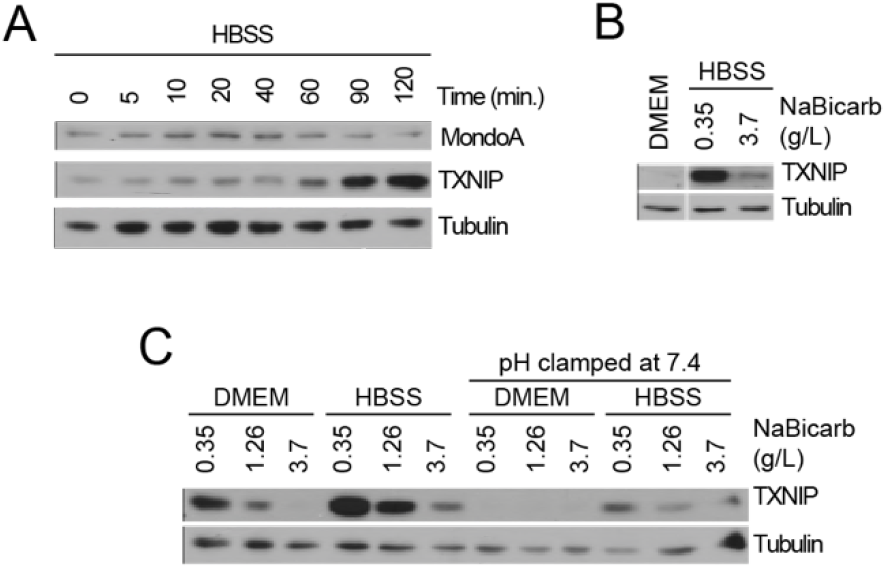
Acidosis drives MondoA transcriptional activity. (**A**) TXNIP and MondoA protein levels in MEFs treated with HBSS as determined immunoblotting. (**B**) TXNIP protein levels of MEFs treated with DMEM, HBSS and HBSS supplemented with sodium bicarbonate to the same amount as DMEM (3.7 g/L). (**C**) TXNIP protein levels in MEFs treated with DMEM and HBSS containing the indicated amounts of sodium bicarbonate. Additionally, the same treatments were clamped to pH 7.4 by adding 25mM HEPES and adjusting the pH with NaOH. ***p<0.001

**Figure 2 – figure supplement 1.**
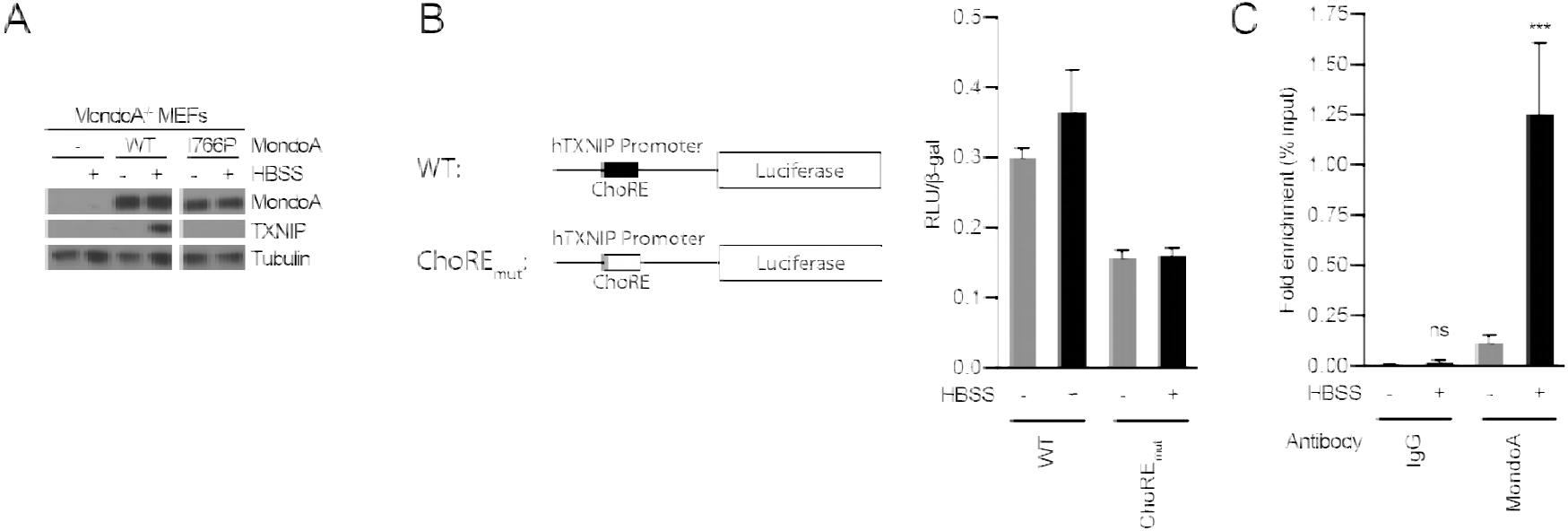
Acidosis drives MondoA transcriptional activity. (**A**) Immunoblot examining TXNIP induction in response to HBSS in MondoA-knockout MEFs complemented with empty vector, wild-type MondoA or MondoA(I766P). (**B**) Schematic depicting the TXNIP-promoter luciferase reporter constructs. A construct was made that contained a mutation in the carbohydrate-responsive element (ChoRE_mut_). Luciferase constructs were transfected into MEFs and HBSS treatment results in a slight induction of luciferase. Using the ChoRE_mut_ TXNIP promoter, initial luciferase expression was lower and HBSS treatment had no effect on luciferase. (**C**) Chromatin-immunoprecipitation performed on MEFs treated with HBSS. Antibodies against MondoA and IgG were used. ***p<0.001; ns – not significant

**Figure 3 – table supplement 1. Differentially regulated genes and overrepresentation analysis (related to Figure 2)**

**Figure 3 – table supplement 2. Enriched gene sets from GSEA (related to Figure 2)**

**Figure 4 – figure supplement 1.**
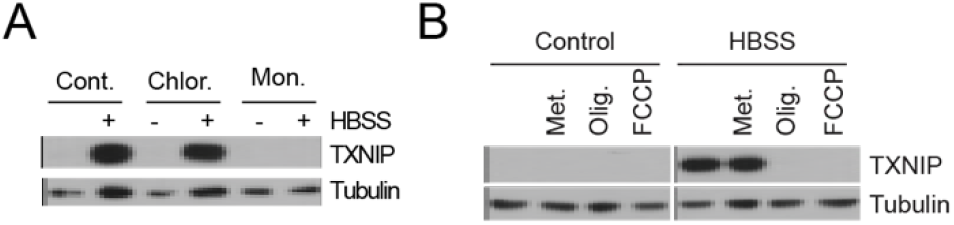
Acidosis-driven MondoA transcriptional activity requires mitochondrial ATP production. (**A**) TXNIP protein levels in MEFs treated with ionophores chloroquine (Chlor., 25 μM) and monensin (Mon., 5 μM), which cause lysosomal and cytosolic alkalization, respectively. (**B**) TXNIP protein levels in MEFs treated with HBSS and the mitochondrial ionophore FCCP or the ETC complex inhibitors metformin (Met., 1 mM) and oligomycin (Olig., 1 μM).

**Figure 5 – figure supplement 1.**
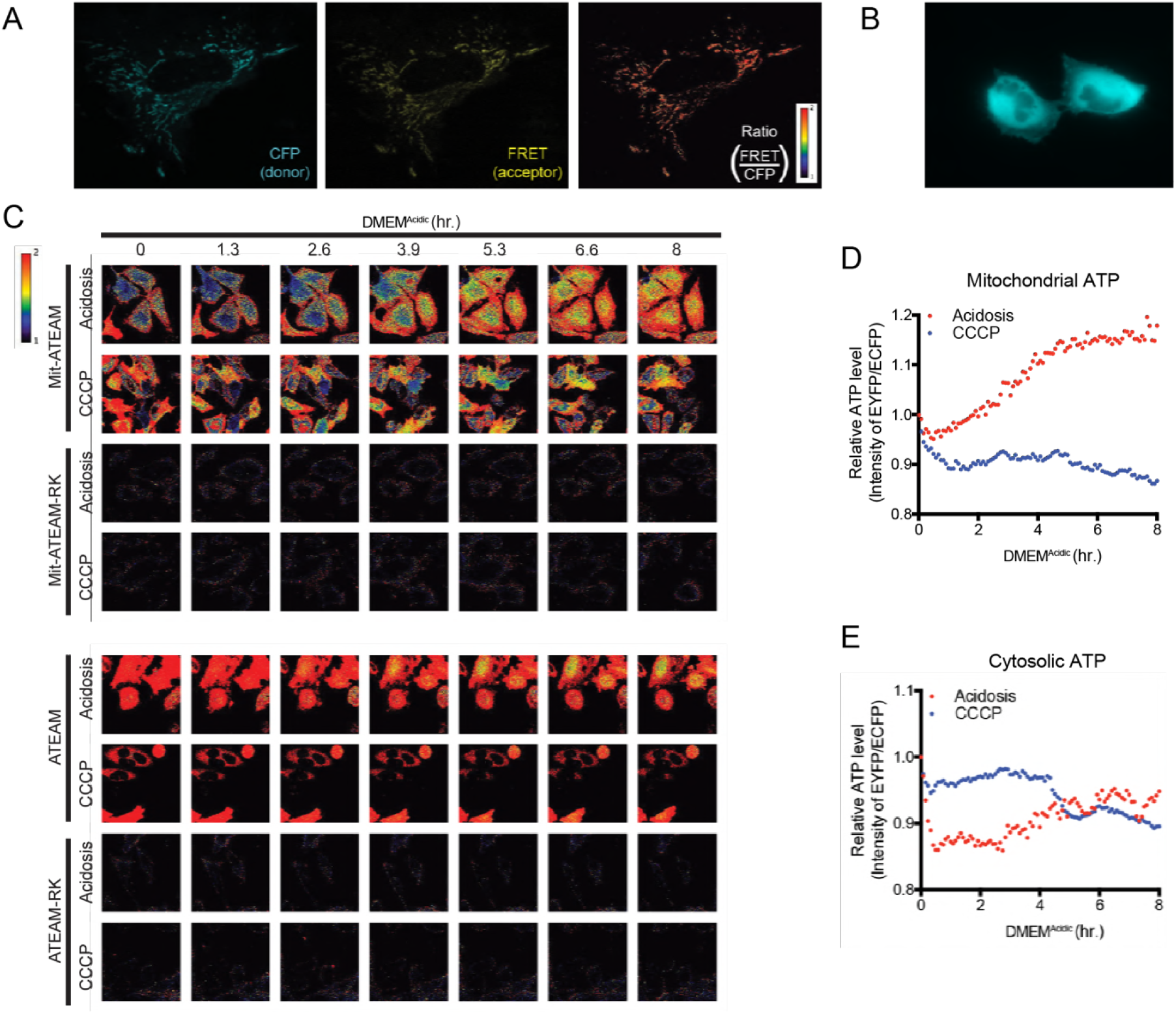
Acidosis drives synthesis of mitochondrial ATP. (**A**) Confocal images at 60X of Mit-ATEAM expressed in HeLa cells. Shown are the CFP and FRET channels as well as the ratio of FRET to CFP (indicating ATP). (**B**) Widefield image at 60X of ATEAM. CFP channel only is shown. (**C**) Confocal images of Mit-ATEAM and Mit-ATEAM(R122K/R126K) in HeLa cells. Cells were treated with DMEM^Acidic^ or CCCP (1 μM) for 8 hours. Images are pseudo-colored according to the FRET/CFP ratio. Notably, the FRET/CFP ratios for Mit-ATEAM(R122K/R126K) and ATEAM(R122K/R126K) was negligible compared to non-mutated constructs. Quantification of (**D**) mitochondrial ATP and (**E**) cytosolic ATP. Confocal images of ATEAM and ATEAM(R122K/R126K) in HeLa cells.

